# Microbiota-dependent early life programming of gastrointestinal motility

**DOI:** 10.1101/2023.11.08.566304

**Authors:** Mary E. Frith, Purna C. Kashyap, David R. Linden, Betty Theriault, Eugene B. Chang

**Affiliations:** Interdisciplinary Scientist Training Program, University of Chicago, Chicago, IL 60637, USA; Department of Medicine, University of Chicago, Chicago, IL 60637, USA; Division of Gastroenterology and Hepatology, Department of Medicine, Mayo Clinic, Rochester, MN 55905, USA; Enteric Neuroscience Program, Department of Physiology and Biomedical Engineering, Mayo Clinic, Rochester, MN 55905, USA; Department of Surgery, University of Chicago, Chicago, IL 60637, USA

**Keywords:** Fecal Microbiota Transplantation, Gastrointestinal Motility, Germ-Free Mice, Postnatal Development, Critical Period, Transit Testing, Whole Gut Transit, RNA Sequencing, Enteric Nervous System, Neurodevelopment

## Abstract

Gastrointestinal microbes modulate peristalsis and stimulate the enteric nervous system (ENS), whose development, as in the central nervous system (CNS), continues into the murine postweaning period. Given that adult CNS function depends on stimuli received during critical periods of postnatal development, we hypothesized that adult ENS function, namely motility, depends on microbial stimuli during similar critical periods. We gave fecal microbiota transplantation (FMT) to germ-free mice at weaning or as adults and found that only the mice given FMT at weaning recovered normal transit, while those given FMT as adults showed limited improvements. RNAseq of colonic muscularis propria revealed enrichments in neuron developmental pathways in mice exposed to gut microbes earlier in life, while mice exposed later – or not at all – showed exaggerated expression of inflammatory pathways. These findings highlight a microbiota-dependent sensitive period in ENS development, pointing to potential roles of the early life microbiome in later life dysmotility.

## Introduction

Early life represents a critical time for host-microbe interactions in the colon, where enteric nervous system (ENS) and immune development continue into post-weaning life. Early life microbial deprivation irreversibly impairs immune development,^1^ but what long term effects does early life microbial deprivation have on ENS development and its functional output, gastrointestinal motility? Given the existence of stimuli-dependent critical periods in central nervous system (CNS) development and motor function,^2,3^ it seems likely that there exist microbial stimuli-dependent critical periods for ENS development and gastrointestinal motility.

Results from several studies support a microbiota-dependent critical period for motility, while not testing it directly. One study in 2014 showed that neonatal mice lacking microbes (“germ-free”, GF) had reduced neuronal fiber density and altered intestinal contractility at postnatal day 3 (P3).^4^ However, because these mice were not followed to adulthood, the extent to which those abnormalities were recoverable was not assessed. Another study identified a potential mechanism by which microbes affect ENS postnatal development involving microbiota-dependent BMP4 and CSF1 signaling between muscularis macrophages and neurons.^5^ However, this study did not assess adult mice, and therefore the long-term consequences of microbiota on ENS maturation were not evaluated. Another study showed that the effect of antibiotic treatment on intestinal serotonin production was more pronounced when given in early adulthood (P42) than when given at birth (P0) or weaning (P21). However, early life was not the focus of the study and these differences were not explored in the context of long term motility changes.^6^ Another study reported that microbial colonization of adult GF mice increased serotonin receptor 5HT4 immunoreactivity and decreased the proportion of Nestin+ to HuCD+ cells in colonic myenteric plexus, but the study did not assess early versus later life microbial exposure.^7^ Therefore, the question has remained about the extent to which early life is a microbiota-dependent critical period in ENS development.

We hypothesized that 1) depletion of gut microbes during a critical window of postnatal development impairs adult GI motility, and 2) this is mediated by changes in neurodevelopmental pathways that impede ENS maturation.

We devised an approach in which germ-free mice were conventionalized (given fecal microbiota transplantation, FMT) with SPF microbes at weaning or as adults; this approach accounted both for time since FMT and age of testing. We found that FMT of GF mice at weaning, but not adulthood, restored normal transit times. These functional changes were accompanied by a normalization in colonic muscularis propria of immune and Hedgehog signaling-dependent genes and pathways in early conventionalized mice, while these remained impaired in late-conventionalized and GF mice. Similarly, despite receiving FMT from the same source, late conventionalized mice developed a microbiota that was less diverse and more divergent from SPF compared to that of early conventionalized mice. These microbiota changes were independent of time since FMT and suggest that the host side developmental changes due to lacking a microbiota earlier in life limited engraftment of the SPF consortium. Finally, in line with lower expression of Hedgehog pathway related transcriptional programs (of which a primary function is to maintain the glial population by inhibiting terminal differentiation of progenitor cells into neurons),^8,9^ the colonic myenteric plexus of late conventionalized and GF mice had a lower glia to neuron ratio.

Our findings highlight time-sensitive ENS-microbiome interactions during development, a concept with notable clinical implications. Motility disorders such as irritable bowel syndrome (IBS) and functional constipation are usually first diagnosed and then managed in adulthood,^10,11^ but earlier intervention or preventative measures during childhood may be more effective toward mitigating lifelong dysmotility.

## Results

### The Early Life Microbiome Affects GI Transit Later in Life

To test whether the presence of microbes during early life is critical for long-term motility, female C57Bl/6 GF were conventionalized using fecal microbiota transplantation (FMT) with pooled SPF fecal slurry at weaning (3-4 weeks [w] of age), 8w, or 12w of age. Female mice were used given the higher prevalence of functional bowel disorders in females.^12^ After 4w (to allow for stabilization of the microbiota) or at 16w of age, we tested the transit times of the mice as a readout of ENS function. This study design (Figure 1A) accounted for potential confounders of age of testing and time post conventionalization. We found that transit times were restored in mice given FMT at weaning (4wC) both 4w after FMT and at 16w of age, while transit was significantly delayed for mice given FMT as adults (8wC, 12wC) (Fig. 1B). The transit deficit was not explained by differences in fecal water content or colon length (Fig. S1). These findings suggests that the critical window for ENS-microbiome interactions for the development of normal motility closed before adulthood.

**Figure 1.**
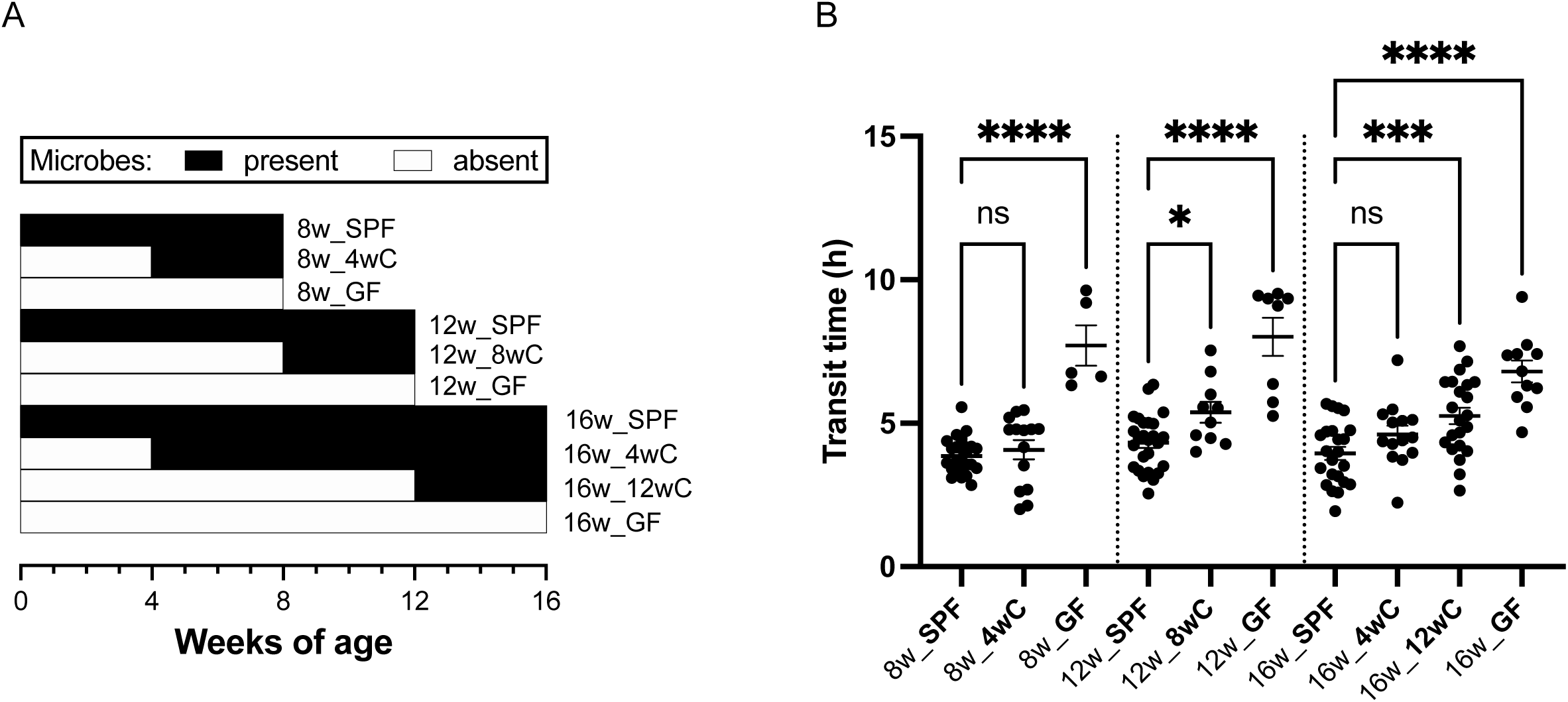
The early life microbiome affects GI transit later in life. (A) Diagram of experimental design. Mice were given FMT at different ages and compared to age matched SPF and GF controls either 4 weeks after FMT or at 16w of age. This design controlled both for time since FMT and age at testing. (B) Mice that lacked a microbiota before adulthood (GF, 8wC, and 12wC mice) failed to recover normal transit times after FMT. Mice given FMT at weaning (4wC) had normal transit after the introduction of microbes. Ordinary one-way ANOVA with Holmes-Sidak multiple comparisons test. All mice shown in the figure are female C57Bl/6. Mean +/- SEM indicated. *p<0.05, ***p<0.001, ****p<0.0001. ns=not significant. See also Figure S1.

### The Early Life Microbiome Affects Adult Gene Expression in Colonic ENS

To explore whether the early life microbiome affects gene expression in the adult ENS, we performed RNA sequencing (RNAseq) of colonic muscularis propria (MP) of adult SPF, GF, 4wC and 12wC mice. We found that each of the groups had distinct signatures of gene expression, indicating that the post-weaning microbiome affects adult gene expression patterns in colonic ENS (Fig. 2). However, the early conventionalized mice appeared most like SPF, as 4wC had almost an order of magnitude fewer differentially expressed (DE) genes than 12wC when compared with SPF (166 versus 1604 genes with padj < 0.05). A PCA plot of colonic MP also shows 4wC as the group closest on average to SPF along PC1 and PC2, while still being distinct from SPF (Fig. 2). This suggests a gradient of influence of the microbiome on postnatal ENS development for which earlier restoration of microbial signals facilitates greater normalization of transcriptional programming.

**Figure 2.**
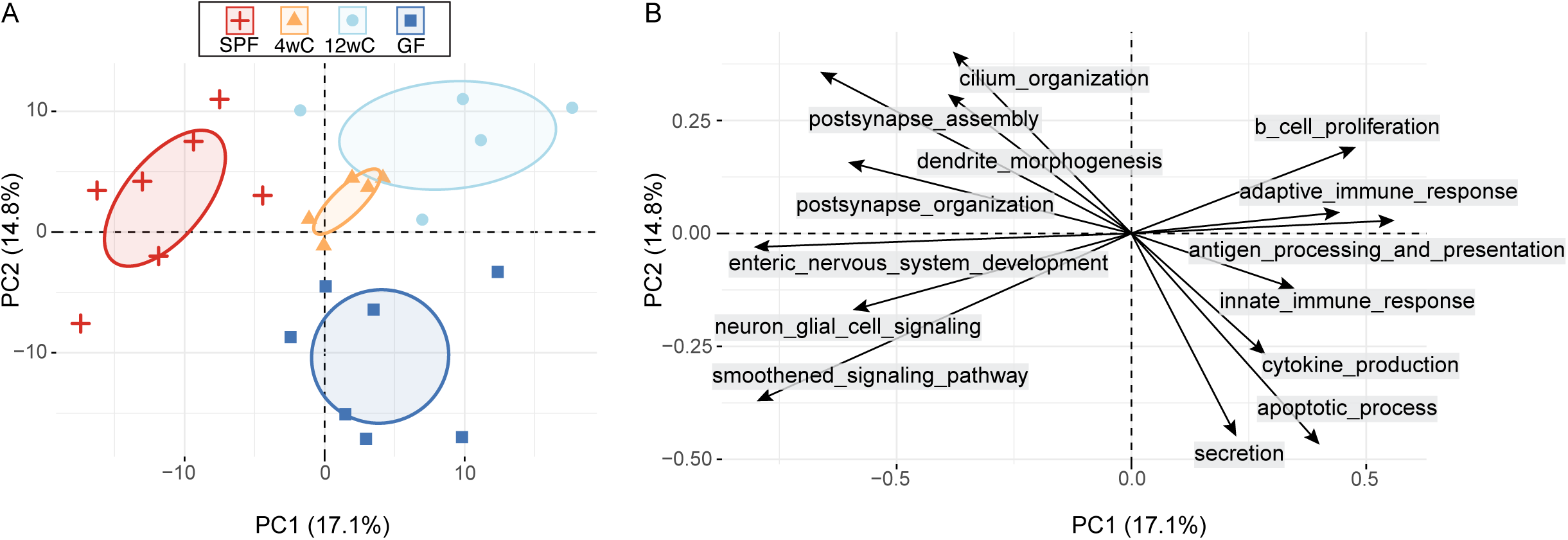
The early life microbiome affects adult gene expression patterns in colonic ENS. (A) PCA plot of gene expression in colonic muscularis propria shows 4wC as the group closest on average to SPF, while retaining similarities to 12wC and GF. (B) GOBP pathway projections onto the dimensions shown in (A). The differences in gene expression between groups can be largely accounted for by higher expression of key neurodevelopmental pathways in SPF and 4wC and higher expression of immune related pathways in GF and 12wC. Note that the axis ranges are different in (A) and (B), such that (B) is a “zoomed in” version of (A); a single pathway (or gene set) in isolation does not explain the group differences but rather the combined contribution of related pathways. Arrow direction represents the relative correlation between the pathway and each component, and arrow length represents the predicted contribution of the pathway to the component (based on expression of genes in that gene set across individuals). See also Figure S2.

To investigate early life microbiota-dependent transcriptional programs in ENS, we compared the DE pathways between groups that either had (SPF, 4wC) or lacked (GF, 12wC) an early life microbiome. Shared upregulated pathways represent cellular programs that rely on microbial signals during a critical window between weaning and adulthood, while shared downregulated pathways represent cellular programs normally kept in check by microbial exposure before adulthood. Two major themes emerged (Figure 2B). In SPF and 4wC, shared upregulated pathways (relative to GF and 12wC, respectively) coalesced on functions or pathways downstream of Hedgehog (Hh) signaling. These included the Smoothened pathway, which is critical to enteric nervous system development and takes place in the neuronal primary cilium (elements which were each represented among the enriched pathways and genes). Importantly, Smoothened signaling promotes maintenance of a robust glial population by inhibiting premature terminal differentiation of neural precursor cells into neurons.^8,9^ The second theme, represented by the shared downregulated pathways in SPF and 4wC, was overwhelmingly immune related (both innate and adaptive). The higher expression of immune pathways in GF and 12wC may reflect an elevated basal level of colonic inflammation in the absence of microbes in GF and an exaggerated immune response to commensal microbes in 12wC. These results strongly suggested that Hedgehog pathway signaling and immune tolerance in colonic ENS rely on microbial signals between weaning and adulthood.

To identify specific genes whose expression depends most strongly on early life microbial signals, we overlapped lists of shared DE genes (p<0.05) up- and down-regulated in SPF and 4wC, each compared to 12wC and GF (Figure S2). The resulting two lists represent genes whose expression most strongly either increased to SPF’s normal levels from GF’s low levels (“recovered-up”; 57 genes) or decreased to SPF’s normal levels from GF’s high levels (“recovered-down”; 95 genes) in 4wC only (Fig. S2A, Table S1). Individual expression levels of these top early life microbiota-dependent genes are plotted as a heatmap in Figure S2B. Like the pathway enrichments, many of these genes relate directly or indirectly to Hh pathway and inflammatory signaling. Thus, we identified genes whose expression in ENS depends on microbiota-originating signals during a critical developmental window.

### Glia-Neuron Ratio Depends on the Early Life Microbiome

Given the Hh pathway’s role in maintaining the pool of glia and neural progenitor cells by inhibiting terminal differentiation into neurons,^8,9^ we wondered whether GF and/or 12wC mice had deficient glia:neuron ratios relative to SPF and 4wC. To explore this possibility, we labeled wholemount preparations of distal colonic muscularis propria with neuronal soma marker Elavl3/4 (HuCD) and glial marker GFAP and quantified the number of neurons and glia per myenteric ganglion. Indeed, we found that GF and 12wC had reduced glia:neuron ratios compared to SPF, while this ratio was restored in 4wC (Figure 3, Figure S3). Corroborating the immunofluorescent quantification, RNA expression ratios of *Gfap* to the average of *Elavl3* and *Elavl4* were higher in mice that were exposed to microbes during early life (SPF, 4wC) than those not exposed during early life (GF, 12wC) (Fig. 3). This finding is concordant with the upregulation of Smoothened signaling in SPF and 4wC (Fig. 2). Therefore, exposure to the microbiota before adulthood preserved the balance between neurons and their critical support cells in colonic ENS.

**Figure 3.**
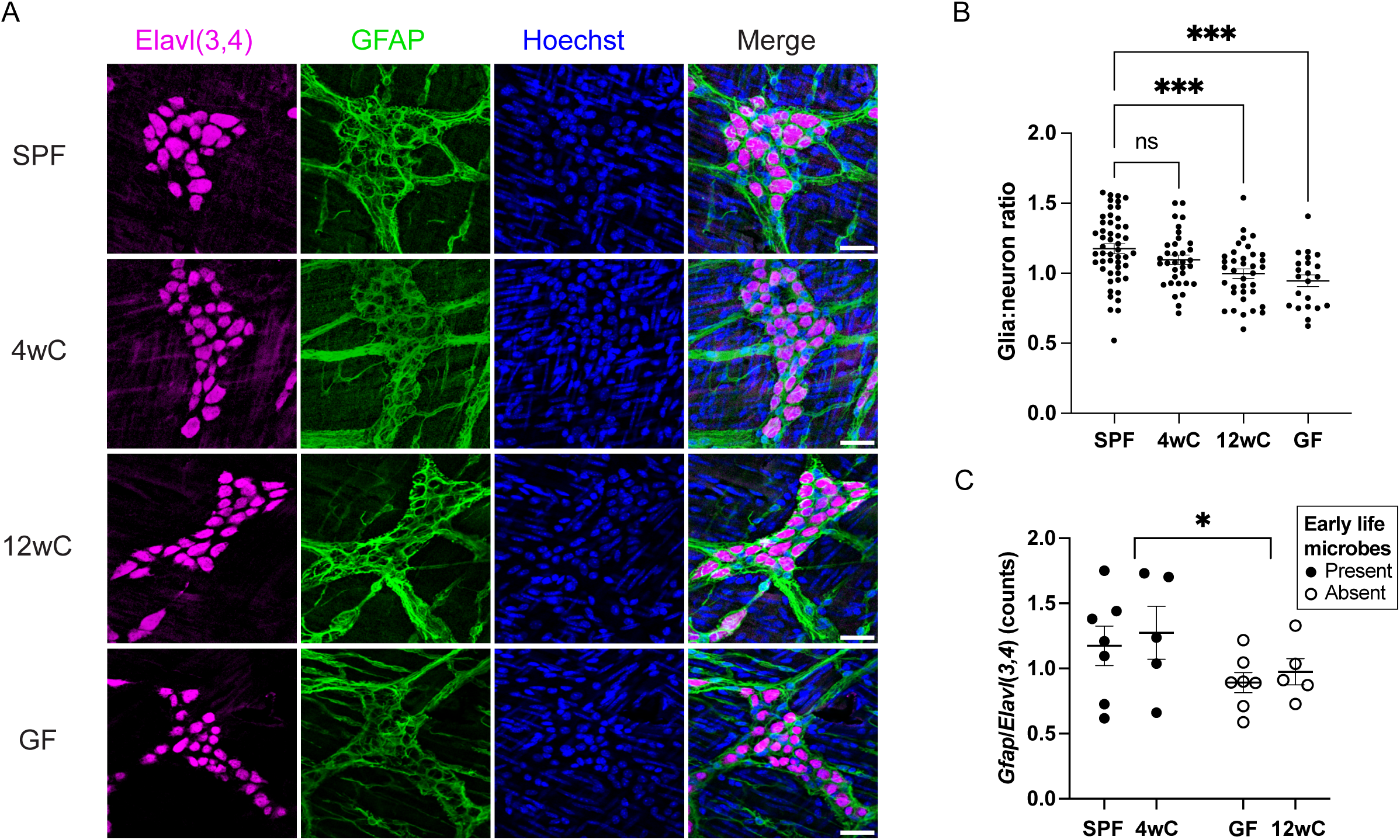
Glia:neuron ratio depends on the early life microbiome. 4wC regains a normal glia:neuron ratio, while 12wC and GF have a lower glia:neuron ratio compared to SPF. (A) Whole-mount immunofluorescent (IF) staining of myenteric plexus in colonic muscularis propria. Representative ganglia from SPF, 4wC, 12wC, and GF, stained for Elavl3/4 (HuCD; neuronal soma), GFAP (glia), and Hoechst (nuclei). Scale bar: 36µm. (B) Ratio of glia to neurons from IF quantification. Data are plotted by ganglion analyzed and include images from 3-6 animals per condition collected over 4 staining batches that each included samples from all conditions. Ordinary one-way ANOVA with Dunnett’s multiple comparisons test. (C) Validation of IF quantification; RNA expression ratio of *Gfap* to the averaged counts of *Elavl3* and *Elavl4* (to correspond to the Elavl(3,4) antibody, which labels both proteins). Comparison is between mice for which microbes were either present or absent during the critical window (i.e., SPF and 4wC for “present”, GF and 12wC for “absent”). Two-way ANOVA. Mean +/- SEM indicated. *p<0.05, ***p<0.001, ns=not significant. See also Figure S3.

We wondered whether the early life microbiota would affect adult neuronal fiber density or neuron subtype distribution in the colonic myenteric plexus. We found no differences in neuronal fiber density in the colonic myenteric plexus of SPF, conventionalized, or GF mice (Figure S4A,B). Concordantly, we found no differences between groups in the RNA expression of general neuronal marker *Tubb3* (Fig. S4C). Notably, there were no neuronal subtype-specific markers whose expression pattern among groups resembled early life microbiome-dependent effects (Fig. S2, Table S1). Taken together, these results suggest that the early life postweaning microbiome regulates the glia:neuron balance and signaling programs of those cells, rather than the specification of neuronal subtypes.

### Adult Microbial Diversity Depends on the Early Life Microbiome

We were interested in whether the fecal microbiota differed in composition or diversity between early and late conventionalized mice once the communities stabilized. Since all conventionalized mice received FMT from the same (SPF-derived) source, such differences would suggest that host-intrinsic physiological factors (based on timing of microbial exposure) shaped the microbiota. Indeed, 16S rRNA amplicon sequencing revealed that the 12wC microbiota diverged in composition from and had lower alpha diversity than 4wC and SPF mice (Figure 4). This was unsurprising, as 12wC’s upregulation of inflammatory pathways in colonic muscularis propria would probably be reflected in increased antimicrobial activity within the colonic mucosa, thereby counteracting colonization by certain commensal microbes. Interestingly, the composition of the FMT gavage solution appears most similar to SPF and 4wC fecal composition when plotted as a PCoA (Fig. 4A), while 12wC shows less overlap. Indeed, 12wC’s sample distances to SPF (weighted Unifrac) were higher than the sample distances between 4wC and SPF (Fig. 4B). Shannon entropy (alpha diversity) was also lower in 12wC (Fig. 4C). This pattern of dissimilarity of 12wC versus similarity of 4wC and SPF suggests that the SPF consortium engrafted more effectively when administered to GF mice at weaning rather than in adulthood. Thus, the immature intestine may be better able to accommodate a normal consortium of microbes than the intestine that has matured without the corresponding microbial stimuli.

**Figure 4.**
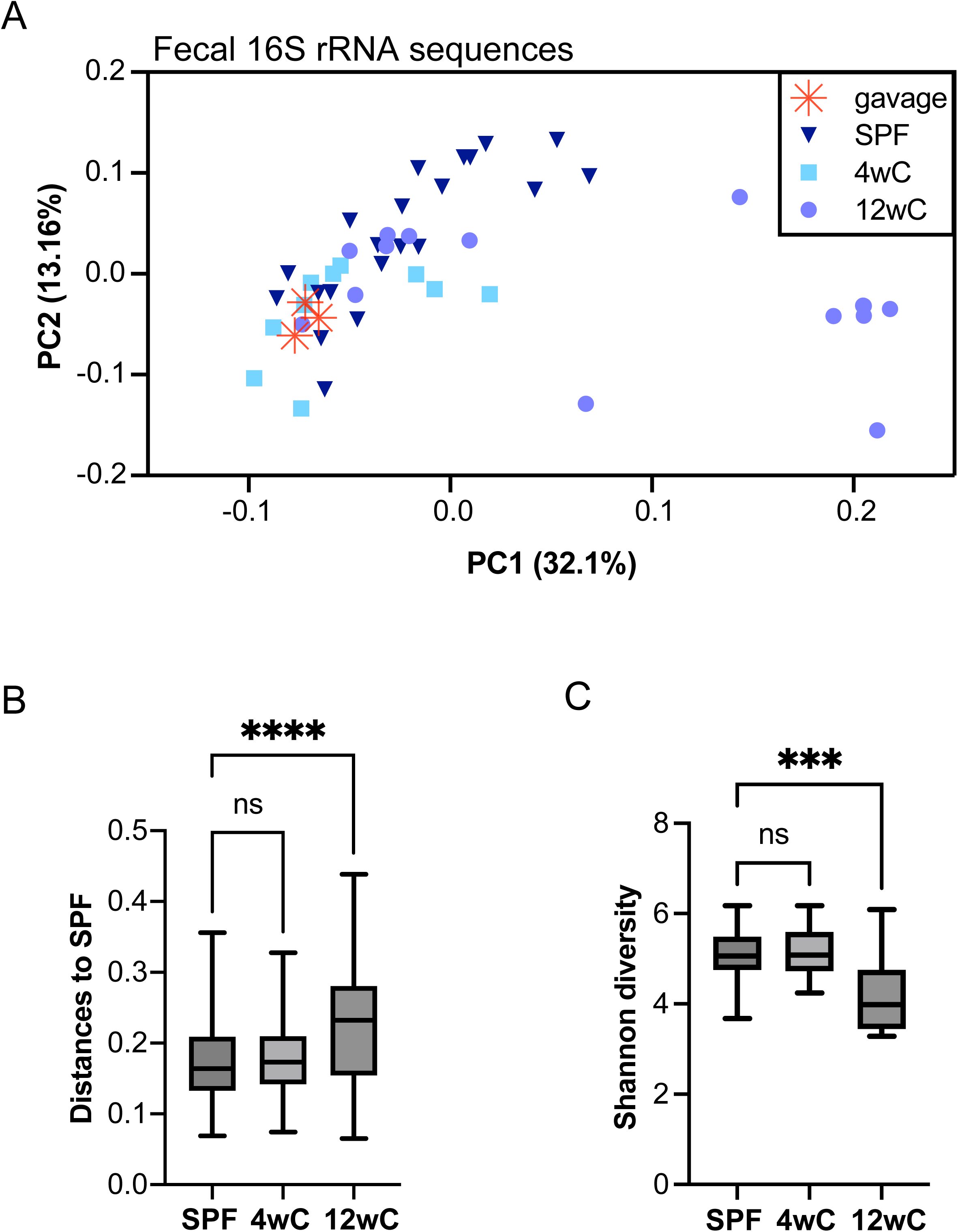
Adult microbial diversity depends on the early life microbiome. (A) Weighted Unifrac PCoA of fecal samples from SPF, 4wC, 12wC, and from aliquots of the gavage solution used for FMT. Note that SPF and 4wC appear more like the gavage solution, while 12wC is more scattered. (B) Weighted Unifrac sample distances to SPF for SPF, 4wC, and 12wC shows that the composition of the 12wC fecal microbiota differed significantly from SPF, while 4wC resembled SPF. Kruskal-Wallis with Dunn’s multiple comparisons test. (C) Shannon entropy scores of microbial alpha diversity; 12wC had lower fecal alpha diversity than SPF, while 4wC did not differ from SPF. One-way ANOVA with Dunnett’s multiple comparisons test. SPF n=21, 4wC n=10, 12wC n=15 samples. Samples collected at 16w of age. Box plot whiskers indicate min to max. ***p<0.001, ****p<0.0001, ns=not significant.

In summary, we have identified a critical window for the development of normal motility that requires the microbiome. The post-weaning microbiome affects adult gene expression patterns in colonic ENS, and earlier restoration of microbial signals facilitates greater normalization of cellular programming related to Hedgehog pathway and inflammatory signaling. We identified a set of genes whose expression in colonic ENS depends on microbiota-originating signals during a critical developmental window. We found that the life microbiome regulates the glia:neuron balance and gene expression profiles, rather than the specification of neuronal subtypes. Finally, we showed that the immature intestine accommodates a new consortium of microbes more fully than the mature intestine.

## Discussion

Fundamental to our understanding of human development is that we need certain experiences during early life to develop normally as adults – this shapes public health guidelines for children. However, while such critical periods in brain development are well accepted, less is known about whether other parts of the nervous system - such as the ENS - are similarly impacted by early life stimuli.

To explore the possibility of a microbiota-dependent critical period for postnatal ENS development and motility, we exposed female germ-free (GF) mice to fecal microbes (FMT) at the time of weaning (4wC) or as adults (12wC) and compared them to age matched GF and conventionally raised (SPF) controls. We found that only the mice given FMT during early life developed normal motility, while those given FMT as adults remained impaired. The transit differences were not explained by differences in fecal water content, colon length, time since FMT, or age of testing. These findings revealed a microbiota-dependent critical period in ENS development affecting adult motility.

Probing potential mechanisms of the motility differences, we performed RNA-sequencing of colonic muscularis propria from adult SPF, 4wC, 12wC, and GF mice. We found that expression of genes related to Hedgehog (Hh) signaling that had low expression in GF mice increased to normal levels in 4wC, but not 12wC, mice. Additionally, we found that expression of genes related to widespread immune activation that were highly expressed in GF mice were also highly expressed in 12wC, while 4wC’s expression normalized to those of SPF; similar immune effects have been reported elsewhere.^1^ Beyond replicating previous results, though, the present work revealed that a “setpoint” for Hh pathway expression in colonic ENS is microbe-dependent and established before adulthood.

One of the functions of the Hh pathway is to maintain a pool of neural progenitor cells by inhibiting premature differentiation into neurons.^13^ We found that 12wC and GF mice had lower glia:neuron ratios than SPF controls, while 4wC ratios did not differ from SPF. Concordantly, a previous study found that deficient Hh signaling in a mouse model of Hirschsprung disease decreased the ratio of glia to neurons in the ENS.^14^ Demonstrating the capacity of the microbiota to activate Hh signaling, another study found that two microbial populations differentially regulated Hh signaling in female mice.^15^ Taken together, the findings of the present study propose a potential mechanistic link between the early life microbiota, Hh pathway regulation, and gastrointestinal motility.

Indeed, several of the developmental phenomena normally occurring around the time of weaning relate to Hh signaling. These include neuron differentiation, refinement of neuronal connectivity, and pathfinding by neuron projections.^8,16–19^ These processes participate in critical period closure in the brain^20–23^ and could have a similar role in the closure of a microbiome-sensitive critical period in ENS. For example, the disrupted glia:neuron balance in mice that lacked microbes during early life could represent a similar process of reduction of differentiation potential of neuron progenitors that characterizes critical period closure in the brain.^21^ Similarly, refinement of neuronal connections is a glia-assisted process that peaks shortly after weaning in rodents^17,24^ and involves phagocytosis and/or autophagy of synapses that have received minimal stimulation.^17,25–27^ Appropriately, several of the early life microbiota-dependent genes in the present study relate to autophagy (e.g., *Tsc1*, *Vps13c*, *Herc1*). Thus, enteric neurons that are not stimulated by microbial stimuli before refinement may lose the opportunity to properly respond. Finally, while we found no differences in density of innervation between groups, it is possible that the connections in 12wC and GF are slightly off target. If so, this might explain the motility defects. As such, in considering the involvement of the Hh pathway in each of these critical postweaning developmental processes, future studies can begin to piece together a plausible mechanism for microbiota-dependent critical period in postnatal ENS development.

Of course, it is possible is that the ENS critical period effects are secondary to immune effects, as the elevated inflammatory signaling in 12wC and GF could impact neuronal function. While this cannot be ruled out by the present study, the motility impairment exists regardless of whether the effects of the early life microbiota are primarily immune or neuronal. It is indeed likely that critical period mechanisms in immune and ENS development interact. An example from the present study: one of the recovered upregulated genes, *Tacr2*, expressed on neuronal varicosities, skews T cells toward a more tolerant phenotype^28,29^ and has antiproliferative immune effects.^30^ Moreover, while a microbiota-dependent critical window for immune development has been described,^1,31^ far less is known about such a critical period for ENS, and this study begins to address that gap.

All in all, our findings support a model in which disruption or depletion of gut microbes during a critical window of postnatal development impairs adult GI motility. Time-sensitive and microbe-dependent developmental signaling pathways may not be able to proceed in the absence or lack of certain microbial signals. Compensations may occur that ultimately ‘wire’ or configure the ENS in such a way as to not properly function or respond to other stimuli once the mouse has more fully matured. Among the signaling programs most impacted by lacking gut microbes during postweaning development are those downstream of the Hedgehog pathway and are likely to involve neuroglial communication. While some of these impairments are reversible, others are not, because subsequent developmental events may have built on the earlier ones and as such may have closed the critical window.

One of the canonical principles in psychology and neuroscience is that early life exposure to certain stimuli – or lack thereof – impacts development. This principle has prompted the development of public health metrics such as the Adverse Childhood Experiences (ACE) score to assess health risks in adulthood.^32^ Screening for early life microbiome disturbances may be important as well. While early life studies often focus on the pre-weaning period in mice, development continues after weaning, and disruption during this phase may have distinct health consequences.

Our findings provide insights into the fundamental significance of environmental disturbances during postweaning development, and the relevance of such disturbances for GI health.

## Supporting information

Key Resource Table

## Acknowledgements

The authors thank the UChicago Functional Genomics and Integrated Light Microscopy Core Facilities, Gnotobiotic Research Animal Facility and Animal Resources Center; and Argonne National Laboratory Environmental Sequencing Facility. MF thanks Jason Koval, Vanessa Leone, Yogesh Bhattarai for technical advice and coordination; and Fran Jackson for administrative support.

Research reported in this publication was supported by the UChicago DDRCC, Center for Interdisciplinary Study of Inflammatory Intestinal Disorders (NIDDK P30 DK042086). MF is supported by NIH T32 GM007281 and NIH F30DK126309. DRL is supported by NIH R01DK129315. PCK is supported by NIH R01DK114007. The content is solely the responsibility of the authors and does not necessarily represent the official views of the NIH.

## Author Contributions

MF, EBC, PCK, and DRL conceptualized the study and reviewed and edited the manuscript. MF performed the experiments and collected the data and prepared the original manuscript, including data curation, formal analysis, and visualization. EBC and MF acquired funding. BT provided technical support for execution of the gnotobiotic mouse studies.

### Lead author

Mary E. Frith

### Corresponding author

Eugene B. Chang

### Declaration of Interests

The authors declare no competing interests.

## STAR Methods

### RESOURCE AVAILABILITY

#### Lead contact

Requests for further information should be directed to the lead contact, Eugene B. Chang.

#### Materials availability

This study did not generate new unique reagents.

#### Data and code availability

RNA sequencing data have been deposited at Gene Expression Omnibus. Microbial 16S rRNA gene sequencing data have been deposited at NCBI Sequence Read Archive. Both are publicly available as of the date of publication. Accession numbers are listed in the key resources table. Additional raw data from Figures 1, 3, 4, S1, and S4 were deposited on Mendeley at doi: 10.17632/snpf8f2tcx.1.

This paper does not report original code.

Any additional information required to reanalyze the data reported in this paper is available from the lead contact upon request.

### EXPERIMENTAL MODEL AND STUDY PARTICIPANT DETAILS

#### Animals

All mice in the study were C57Bl/6. Germ-free female C57Bl/6 mice were given FMT at weaning between 3-4w of age (“4wC”) or between 11-13w of age (“12wC”). Female mice were chosen as functional bowel disorders are more prevalent in females than males.^12^ Germ-free and SPF adult mice between 12-25w of age were used as controls for similarly aged conventionalized mice. Age-matched untreated SPF mice were used as controls. Animals were housed in the University of Chicago Animal Resources Center on a 12:12h light:dark schedule under specific pathogen-free or germ-free conditions. All study procedures were approved by the University of Chicago Institutional Animal Care and Use Committee (IACUC).

### METHOD DETAIL

#### Transit testing

Transit testing was performed as previously described.^33^ Mice were placed individually into a cage with bedding removed and a steel wire rack placed on the floor of the cage so that fecal pellets would fall below. A white paper towel was placed underneath each cage to facilitate visualization of the pellets. At the start of testing, the mice were gavaged with 100-300μl 6% carmine dye (Sigma-Aldrich C1022) in 0.5% methyl cellulose (Sigma-Aldrich M0512) and the time was recorded. Whole gut transit time was defined as the interval between the dye gavage and the appearance of a fully red stool pellet, verified by smearing the pellet onto a white paper towel to ensure it was red throughout the pellet. This method has been well validated in conventional mice and there have been no reported adverse events.

#### Fecal microbiota transplantation

Fecal samples from SPF mice between 8-16 weeks of age were collected over the course of several days and stored at −20 deg C until enough had been collected to last through the entire study. The stool samples were pooled, thawed on ice, and mixed thoroughly with a spatula. The pooled stool was divided into 100mg aliquots and sterile PBS was added to a total volume of 1mL. Each aliquot was vortexed for 1 minute with the PBS prior to freezing at −80deg C. On the day of the FMT, the aliquot was thawed, vortexed for 1 minute, then particulate matter was briefly spun down. The supernatant was transferred into a separate tube and this solution was used for oral gavage. For FMT, mice were transferred from the gnotobiotic facility to the barrier mouse facility on the day of the FMT.

#### Tissue harvest

Mice were euthanized by CO_2_ asphyxiation, the abdominal cavity was opened, and mice were transcardially perfused with PBS. Colon and cecum were dissected out on ice. Stool was removed from the colon and the colon was opened lengthwise. Approximately 1.5cm of the proximal and distal ends were collected for immunohistochemistry and pinned flat onto a silicon dish, washed with PBS, then 4 percent PFA in PBS was added while tissue harvest was completed. The remaining portion of the colon was placed into RNALater Stabilization Solution (Invitrogen, AM7024) and stored at 4 deg C. The 4 percent PFA solution was removed at the end of the dissection and freshly made Zamboni’s fixative was added to the tissue, which was gently agitated overnight at 4 deg C, washed 3x in PBS, followed by washing in PBS + 10 percent sucrose solution at 4 deg for several hours, then agitated gently overnight in PBS + 20 percent sucrose + 10 percent glycerol solution, rinsed once in PBS and then stored in PBS + 0.1% sodium azide at 4 deg C.^34^ For both wholemount immunohistochemistry and for RNA extraction, the epithelium was removed from the colonic muscularis propria and only the muscularis propria was used for downstream applications.

#### Immunohistochemistry

For staining, intestinal segments were pinned into another silicon coated dish. Heat induced epitope retrieval (HIER) was performed in citrate buffer at 80-95 deg C for up to 50 min. Following protein block and permeabilization in Superblock buffer (PI37515, Thermo Scientific) with 0.3% TX100 for 30 min at 37 deg C, Mouse-on-mouse blocking was performed with MOM Blocking Reagent (Vector Labs; 5 drops per 9ml of PBS + 0.1% TX100) and incubated at 37 deg C for 1 hour. After washing with PBS + 0.1% TX100 (“PBST”), primary antibody incubation with 1:500 mouse monoclonal HuC/HuD (A21271, Invitrogen) and 1:2000 chicken polyclonal GFAP (ab4674, Abcam) or with mouse monoclonal 1:500 Tubb3/Tuj1 (Biolegend 801202) in Superblock w/ 0.3% TX100 at room temperature (RT) shaking overnight. After 5-7 washes with PBST over the course of 1-3 hours, secondary antibody incubation (Alexa Fluor dyes, Molecular Probes) was performed in Superblock with 0.3% TX100 shielded from light at RT shaking for 1 hour. Nuclear staining was performed with Hoechst 10ug/ml in PBST shielded from light at RT shaking for 10 minutes. 4-5 washes in PBST in 45 minutes then 3 brief PBS washes were performed prior to mounting in Prolong Gold in a glass bottom dish.

#### RNA extraction and sequencing

RNeasy Mini Kit from Qiagen (cat # 74104) was used to extract RNA. For the lysis and homogenation step, Qiagen PowerBead Tubes with Garnet 0.70mm (cat #13123-50) were used with a bead beater for 1 minute at 3450 oscillations/min (GlenMills, Beadbeater-16).

Next generation RNA sequencing was sequenced with Illumina NovaSeq 6000 (Oligo-dT mRNA directional paired end with 50-60M paired end reads/sample) at the University of Chicago Functional Genomics Core Facility.

#### Microbial DNA extraction and 16S rRNA gene sequencing

DNA was extracted from freshly collected fecal pellets using the Qiagen PowerSoil DNeasy Kit (cat. #47016). DNA was sequenced by Argonne National Laboratories Environmental Sample Preparation and Sequencing Facility using Illumina HiSeq2500 with 150bp length reads using primers for the bacterial 16S gene V4 region.

#### Fecal water content

Stool pellets were freshly collected, weighed, and allowed to dry overnight in an oven set to 55 deg C, then reweighed. The water fraction was calculated as 1 minus (dry weight / wet weight).

#### Microscopy

Wholemount immunofluorescence images were taken with a Leica SP8 microscope at the University of Chicago Light Microscopy Core Facility with a 20X oil immersion objective. Areas to image were selected to maximize the number of neuronal cell bodies in each field of view.

### QUANTIFICATION AND STATISTICAL ANALYSIS

#### RNAseq of colonic muscularis propria

Raw files were downloaded from the UChicago Genomics Core server as fastq.qz files, with four files per sample: R1 and R2 each for two flow cells. UChicago Research Computing Center server was used for initial analysis using Python. Reads 1 and 2 for each sample were concatenated by flow cell. Reads were trimmed using Trimmomatic (0.39).^35^ Alignment was performed using STAR (2.7.9a)^36^ and BAM files were sorted by coordinate. Samtools (1.14)^37^ was used for indexing. featureCounts from subread (2.0.1)^38^ was used to generate counts using mm39 (.gtf file obtained from Ensembl.org).

Differential expression analyses were performed for the female colon samples using DESeq2 (1.40.2).^39^ The SVA (3.48)^40^ package identified 3 surrogate variables which were included as technical covariates in DESeq design formula along with Condition (SPF, GF, 4wC, 12wC). For heatmap plotting, counts were normalized with the variance stabilizing transformation (vst) and log fold changes were shrunk with apeglm.^41^ The factoextra^42^ package was used to create the PCA with ellipses and ComplexHeatmaps^43^ was used to create the heatmap. Pathway enrichment was performed with fgsea (1.26.0)^44^ using the Gene Ontology Biological Process gene set (m5.go.bp.v2023.1.Mm.symbols.gmt) from MSigDB.^45–47^

To identify the early life microbiome dependent gene lists (i.e. those that returned to SPF levels in 4wC only), several pairwise DE (pvalue < 0.05) gene lists were intersected and differentiated. Specifically, for the genes that were recovered "down" to normal expression levels in 4wC relative to SPF (and not in 12wC relative to either group), the following intersections and set differences were made: downregulation in SPF versus GF, SPF versus 12wC, and 4wC versus 12wC; upregulation in GF versus 4wC; and no difference between GF versus 12wC nor between SPF vs 4wC. For “recovered - up" genes, the intersections and set differences were as follows: upregulated in SPF versus GF, SPF versus 12wC, and 4wC versus 12wC; downregulated in GF versus 4wC; and no difference between GF versus 12wC nor between SPF vs 4wC.

#### Microbiome analysis

For microbiome analysis, FastQ files were unzipped with gzip for use with qiime2-2022.2.^48^ These were in the format of emp-paired-end-sequences. Demultiplexing and quality filtering was performed with q2-demux and denoising with q2-dada2.^49^ Features were aligned using mafft^50^ and a taxonomy was constructed with fasttree2^51^ and filtered at the sampling depth 11007, which maximized the number of samples and features retained. For taxonomic analysis, the Greengenes classifier was used at 99% sequence identity. The Shannon metric was used for alpha diversity and Unifrac^52^ distances were used for beta diversity analyses.

#### Statistics

Statistical analysis was performed using GraphPad Prism and R/RStudio.^53,54^ Statistics are provided in the figure legends as applicable. Selection of statistical tests (e.g., ANOVA versus nonparametric alternatives) was based on whether statistical assumptions were upheld. Multiple comparisons correction was performed as appropriate (as indicated in figure legends). For the statistics related to pathway and gene enrichments, the default methods in DESeq2^39^ and fgsea^44^ were used.

#### Image analysis

For glia:neuron ratio, Adobe Photoshop “Lasso” and “Count” tools were used to outline ganglia for ROIs and then count HuCD+ cell bodies (neurons) and GFAP+ cell bodies (glia). Specifically, glia were defined as (1) HuCD-nuclei (2) within the border of GFAP staining delineating ganglion (3) not having an elongated nucleus resembling that of a muscle cell.

For neuronal fiber density, Tubb3/Tuj1 staining was quantified per image as percent area of the field of view at plane at the level of the myenteric plexus. Tuj1 images were processed in ImageJ/Fiji^55^ by smoothing with Gaussian blur with sigma = 0.7, subtracting background with a rolling ball radius of 50 pixels, and thresholded such that background staining was not included in the area calculation. See also Figure S3.

## Notes

### Competing Interest Statement

The authors have declared no competing interest.

