## Supplemental Figures and Tables for "Microbiota-dependent early life programming of gastrointestinal motility"

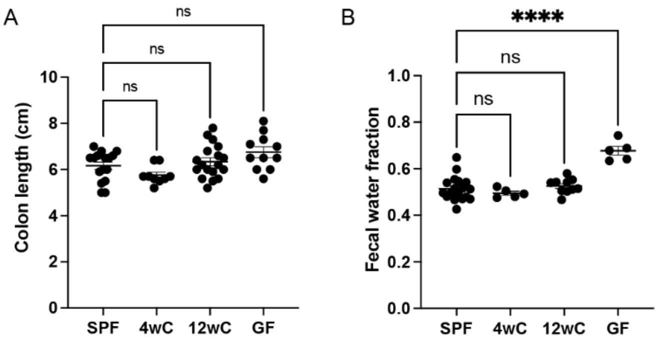


**Figure S1. Slow transit in 12wC is not explained by differences in colon length or fecal water content.**

(A) Colon lengths of 16w-old SPF, 4wC, 12wC, and GF mice did not differ. One-way ANOVA with Dunnett's multiple comparisons correction. (B) Fecal water fraction did not differ between SPF, 4wC, and 12wC, though it was higher in GF. All mice are conventionally raised female C57Bl/6 mice. (A) Colon length: SPF n=16, 4wC n=9, 12wC n=18, GF n=11. (B) Fecal water fraction: SPF n=10, 4wC n=9, 12wC n=10, GF n=5. ns=not significant, ****p<0.0001.

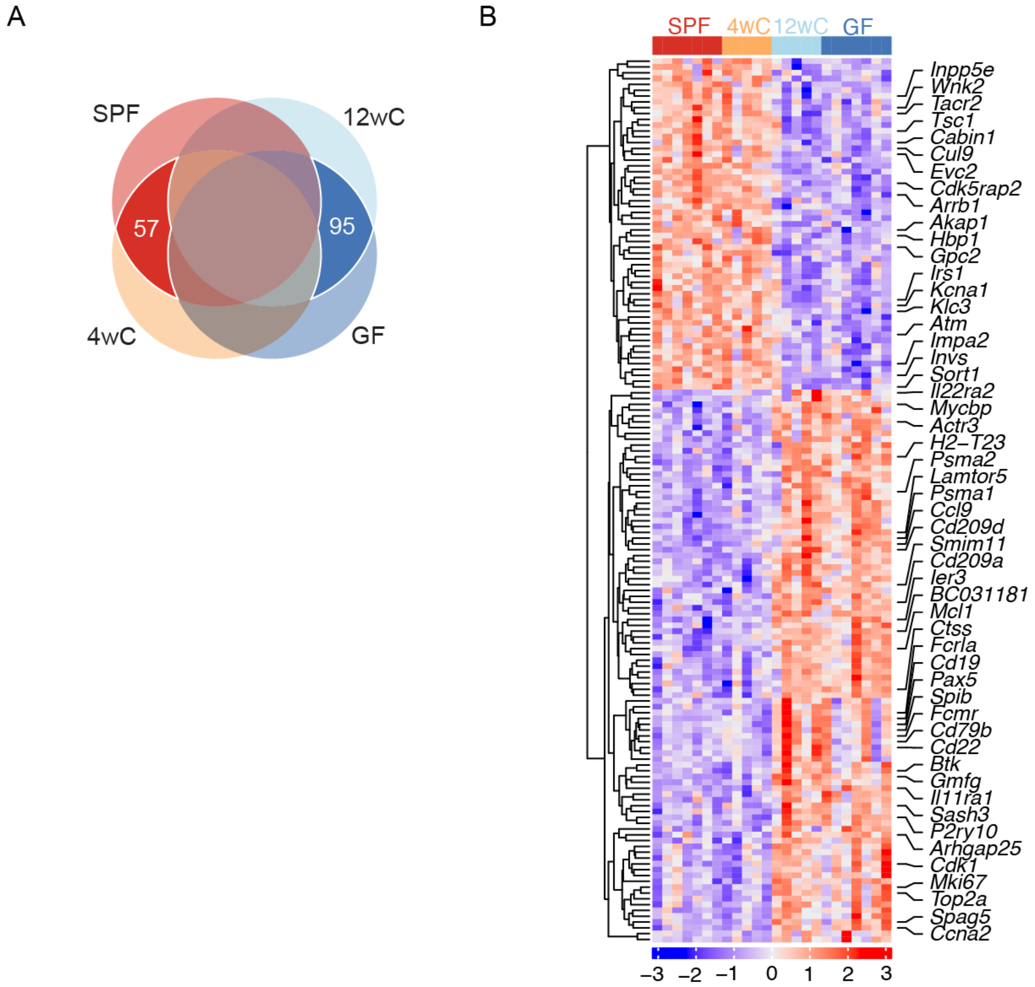


**Figure S2. Early life microbiome-dependent genes.**

(A) Schematic of the group intersections that were made to identify early life microbiome-dependent up- and down-regulated genes. The red section shows the count of genes that were increased in 4wC to SPF's normal expression levels, and the blue section shows the count of genes that were decreased in 4wC to SPF's normal expression levels – that is, genes that remained more highly expressed in GF and 12wC. Differentially expressed ($p<0.05$) gene lists from pairwise group comparisons were intersected and contrasted to identify genes whose normal expression in colonic muscularis propria required the presence of microbes before adulthood. Specifically, for genes that recovered "up" to normal expression levels in 4wC relative to SPF (and not in 12wC relative to either group), the intersections and set differences were as follows: upregulated in (1) SPF versus GF, (2) SPF versus 12wC, and (3) 4wC versus 12wC; downregulated in (4) GF versus 4wC; and no difference between (5) GF versus 12wC nor between (6) SPF vs 4wC. For the genes that were recovered "down" to normal expression levels in 4wC relative to SPF (and not in 12wC relative to either group), the following intersections and set differences were made: downregulation in (1) SPF versus GF, (2) SPF versus 12wC, and (3) 4wC versus 12wC; upregulation in (4) GF versus 4wC; and no difference between (5) GF versus 12wC nor between (6) SPF vs 4wC. (B) Heatmap showing relative expression of the recovered genes across groups, a subset of which are annotated to the right of the heatmap. See Table S1 for the full list of genes. SPF (n=7), GF (n=7), 4wC (n=5), 12wC (n=5).

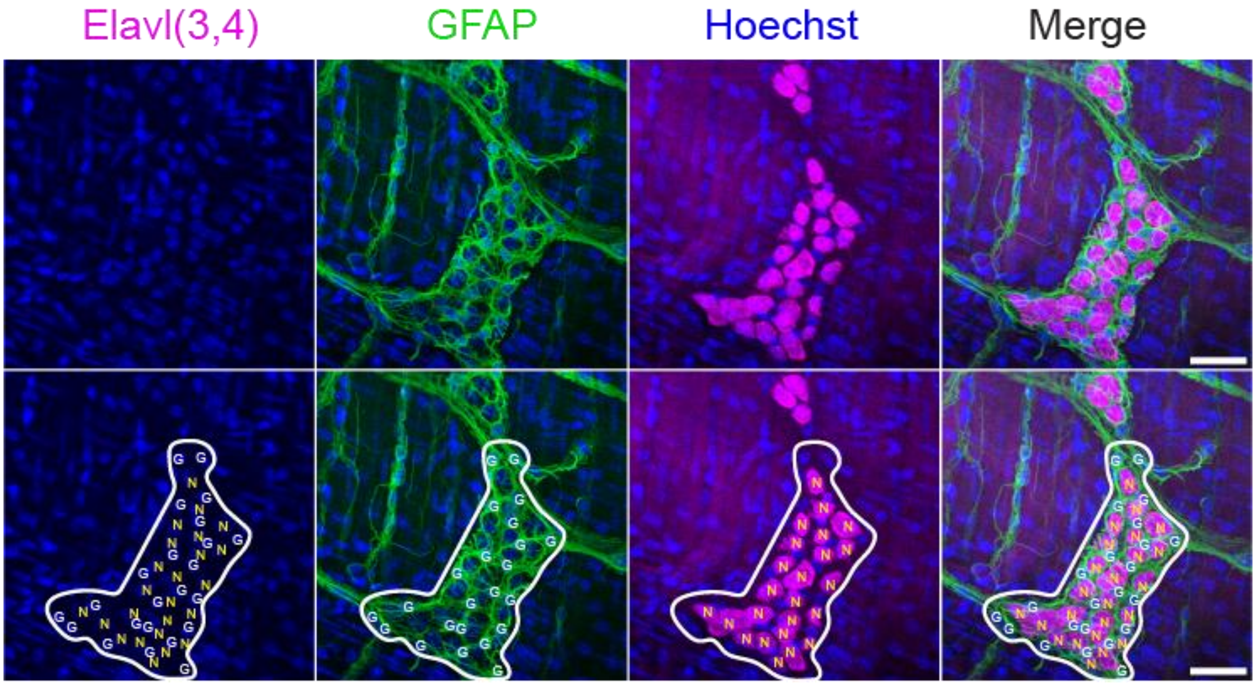


**Figure S3. Example annotation of glia and neurons for analysis of glia:neuron ratio.**

Blue: Hoechst (nuclei); Green: GFAP (glial marker), Magenta: Elavl3/4 (HuC/HuD; neuron soma marker). G=glial cell; N=neuron. Ganglia were defined as a cluster of at least 12 neuronal soma within at most a soma length's distance from the nearest soma (using the length from the longer of the two soma). Nuclear staining aided in determination of whether a given area of marker staining represented one or two cells. Scale bar: 36μm.

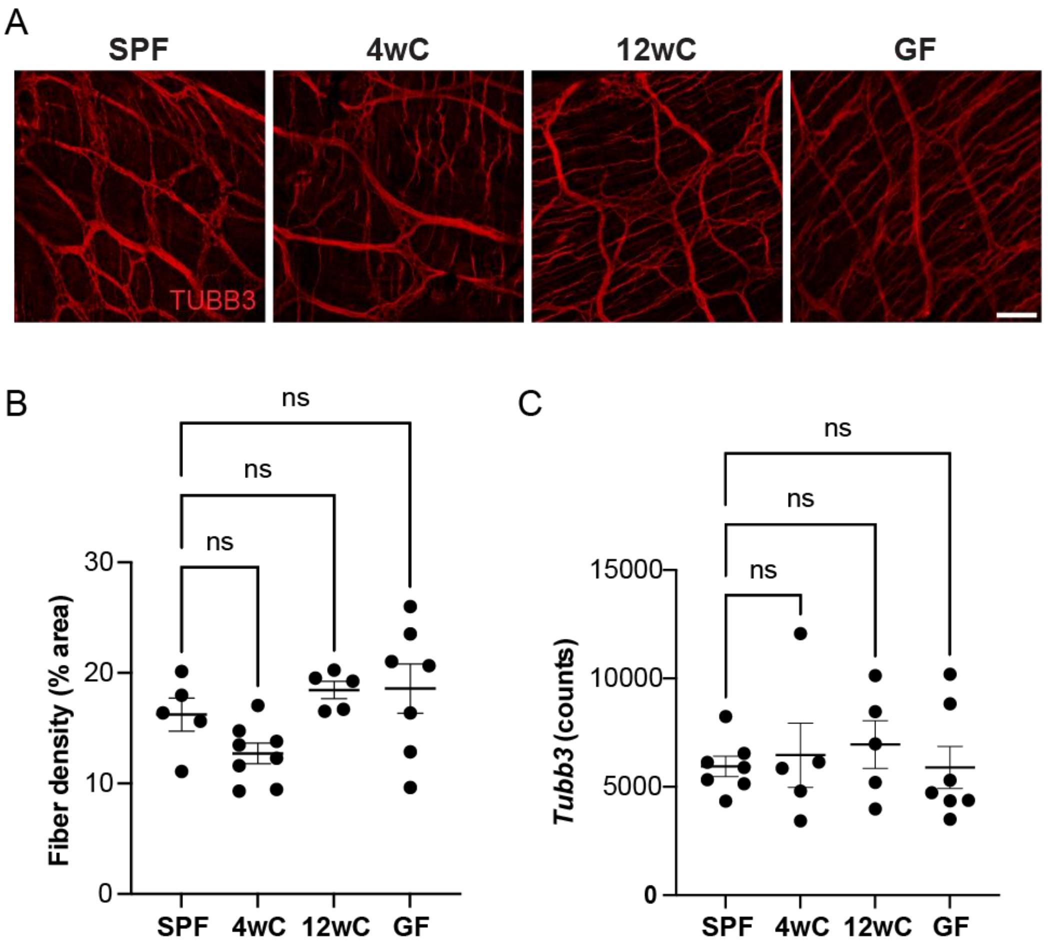


**Figure S4. Neuronal fiber density did not differ based on early life microbiome status.**

(A) Wholemount Tubb3 (Tuj1) immunostaining of distal colon muscularis propria. Representative images from adult female SPF, 4wC, 12wC, and GF mice. Scale bar = 75µm. (B) Neuron fiber density quantification as percent area. Ordinary one-way ANOVA with Dunnett's multiple comparisons test. (C) Tubb3 gene expression as normalized counts. ns=not significant.

Early life microbiota-dependent genes

**Increased** to SPF levels from GF's low levels in 4wC only

| | | | | | |
|---|---|---|---|---|---|
| *Akap1* | *Cnnm3* | *Herc1* | *Krba1* | *Tmod1* | *Zfp882* |
| *Akap8l* | *Cplane1* | *Hsbp1l1* | *Mau2* | *Tsc1* | *Zscan20* |
| *Arrb1* | *Crtc1* | *Hsf4* | *Mroh2a* | *Vps13c* | |
| *Atm* | *Cul9* | *Impa2* | *Nudt13* | *Vwa8* | |
| *Bcl7a* | *Dlg5* | *Inpp5e* | *Nynrin* | *Wnk2* | |
| *Cabin1* | *Erich3* | *Invs* | *Qser1* | *Wrn* | |
| *Ccdc141* | *Evc2* | *Irs1* | *Rad50* | *Yeats2* | |
| *Ccdc77* | *Exd2* | *Kat6b* | *Rgs9* | *Zfp334* | |
| *Ccdc93* | *Gpc2* | *Kcna1* | *Sort1* | *Zfp607a* | |
| *Cdk5rap2* | *H2bc8* | *Klc3* | *Tacr2* | *Zfp677* | |
| *Cep135* | *Hbp1* | *Klhl11* | *Tex15* | *Zfp84* | |

**Decreased** to SPF levels from GF's high levels in 4wC only

| | | | | | |
|---|---|---|---|---|---|
| *AB124611* | *Cd209a* | *Dad1* | *Kif15* | *P2ry10* | *Slc30a5* |
| *Actr3* | *Cd209d* | *Fam241a* | *Knl1* | *Panx1* | *Smim11* |
| *Akr1a1* | *Cd22* | *Fcmr* | *Lamtor5* | *Parl* | *Spag5* |
| *Aldh1a7* | *Cd37* | *Fcrla* | *Ly6d* | *Pax5* | *Spib* |
| *Arf2* | *Cd79b* | *Fis1* | *Mcl1* | *Psma1* | *Taldo1* |
| *Arhgap25* | *Cdca3* | *Gmfg* | *Mki67* | *Psma2* | *Tmem251* |
| *Arpc5* | *Cdk1* | *Gp2* | *Mrpl27* | *Psma6* | *Tmem254* |
| *Atp6v0e* | *Cenpf* | *H2-T23* | *Ms4a1* | *Psmb3* | *Tmem50a* |
| *Atp6v1h* | *Ckap2* | *Hmgn3* | *Mtarc2* | *Psmb5* | *Top2a* |
| *BC031181* | *Clec4a2* | *Hmmr* | *Mycbp* | *Ptpn12* | *Tpx2* |
| *Brip1* | *Crip1* | *Id1* | *Naa12* | *Qdpr* | *Tspan32* |
| *Btk* | *Csrnp1* | *Id3* | *Necap1* | *Rer1* | *Ugcg* |
| *Ccl9* | *Ctsl* | *Ier3* | *Neil3* | *Sash3* | *Vdac3* |
| *Ccna2* | *Ctss* | *Il10ra* | *Nek2* | *Sell* | *Wfdc17* |
| *Ccnb1* | *Cyb5a* | *Il11ra1* | *Oaz1* | *Shcbp1* | *Xlr4a* |
| *Cd19* | *Cycs* | *Il22ra2* | *Orm1* | *Siglecg* | |

**Table S1. Early life microbiome-dependent genes.**

Differentially expressed ($p<0.05$) gene lists from pairwise group comparisons were intersected and differentiated to identify genes whose normal expression in colonic muscularis propria required the presence of microbes before adulthood. See caption of Figure S2, Methods, or the main text for more details.
