## Supplementary material for "Microbiota-dependent early life programming of gastrointestinal motility": Key Resource Table

**KEY RESOURCES TABLE**

| REAGENT or RESOURCE | SOURCE | IDENTIFIER |
| --- | --- | --- |
| Antibodies | | |
| Mouse monoclonal anti-TUBB3 | BioLegend | Cat#801202 |
| Mouse monoclonal anti-HuC/HuD | Invitrogen | Cat#A-21271 |
| Chicken polyclonal anti-GFAP | Abcam | Cat#ab4674 |
| Chemicals, peptides, and recombinant proteins | | |
| Carmine | Sigma-Aldrich | Cat#C1022 |
| Methyl cellulose | Sigma-Aldrich | Cat#M0512 |
| Critical commercial assays | | |
| PowerSoil DNeasy Kit | Qiagen | Cat#47016 |
| RNeasy Mini Kit | Qiagen | Cat#74104 |
| Deposited data | | |
| RNAseq raw fastq data and per sample raw counts | This paper | GEO: GSE241697 |
| Genome annotation for Mus musculus GRCm39 version 109 | Ensembl | https://ftp.ensembl.org/pub/release-109/gtf/mus_musculus/ |
| 16S rRNA amplicon sequencing data | This paper | SRA: PRJNA1021665 |
| Additional raw data for Figs. 1, 3, 4, S1, and S4 | This paper | Mendeley: doi: 10.17632/snpf8f2tcx.1 |
| Experimental models: Organisms/strains | | |
| Mouse: C57Bl/6 | Bred in house | RRID:MGI:2159769 |
| Software and algorithms | | |
| Fiji/ImageJ | Schindelin et al.^1^ | https://fiji.sc/ |
| Photoshop | Adobe | RRID:SCR_014199 |
| DESeq2 | Love et al.^2^ | https://doi.org/doi:10.18129/B9.bioc.DESeq2 |
| STAR | Dobin et al.^3^ | https://github.com/alexdobin/STAR |
| SAMtools | Danecek et al.^4^ | https://www.htslib.org/ |
| featureCounts | Liao et al.^5^ | https://subread.sourceforge.net/ |
| sva | Leek et al.^6^ | https://doi.org/doi:10.18129/B9.bioc.sva |
| RStudio | RStudio Team^7^ | RRID:SCR_000432 |
| R | R Project^8^ | https://www.r-project.org/ |
| GraphPad Prism | GraphPad | RRID:SCR_002798 |
| Qiime2 | Bolyen et al.^9^ | https://qiime2.org/ |
| Unifrac | Lozupone & Knight^10^ | RRID:SCR_014616 |
| Other | | |
| Mouse stool | This study | N/A |
